# Balancing selection on genomic deletion polymorphisms in humans

**DOI:** 10.1101/2022.04.28.489864

**Authors:** Alber Aqil, Leo Speidel, Pavlos Pavlidis, Omer Gokcumen

**Author notes:** **Correspondence:** Omer Gokcumen.

## Abstract

A key question in biology is why genomic variation persists in a population for extended periods. Recent studies have identified examples of genomic deletions that have remained polymorphic in the human lineage for hundreds of millennia, ostensibly owing to balancing selection. Nevertheless, genome-wide investigations of ancient and possibly adaptive deletions remain an imperative exercise. Here, we used simulations to show an excess of ancient allele sharing between modern and archaic human genomes that cannot be explained solely by introgression or ancient structure under neutrality. We identified 63 deletion polymorphisms that emerged before the divergence of humans and Neanderthals and are associated with GWAS traits. We used empirical and simulation-based analyses to show that the haplotypes that harbor these functional ancient deletions have likely been evolving under time- and geography-dependent balancing selection. Collectively, our results suggest that balancing selection may have maintained at least 27% of the functional deletion polymorphisms in humans for hundreds of thousands of years.

## INTRODUCTION

The evolutionary forces that shape the allele frequency distribution of functional genetic variants remain a hotly debated issue. In humans, tens of thousands of common variants have been reported to be associated with human diseases (Loos, 2020). However, the mainstream view remains that the majority of these functional genetic variants have had a negligible effect on reproductive fitness and that the frequency of these variants has fluctuated neutrally by drift over time (Bromberg et al., 2013; Dudley et al., 2012). Functional variants that do have measurable effects on fitness are often observed at a low frequency (Eyre-Walker, 2010). These low-frequency functional variants are considered to be in the process of being eliminated from the population by negative selection (Gibson, 2018; Lettre, 2014; Zeng et al., 2018). Nevertheless, an increasing number of studies are showing that more complex evolutionary histories (Benton et al., 2021; Mathieson & Mathieson, 2018), including introgression from archaic hominins (McArthur et al., 2020), geography-specific adaptation (Hamid et al., 2021; Lachance & Tishkoff, 2013; Mendoza-Revilla et al., 2021), negative selection (Zeng et al., 2018), and polygenic selection (Barghi et al., 2020; Berg & Coop, 2014; Pritchard et al., 2010; Sella & Barton, 2019) may explain the allele frequencies of variants associated with complex diseases. In this context, we aim to test the hypothesis that balancing selection is a considerable force in shaping the allele frequencies of extant functional deletions in the human genome.

Balancing selection is a mode of natural selection that maintains a genomic variant at an intermediate frequency by overcoming its stochastic loss or fixation by genetic drift (Fijarczyk & Babik, 2015; Fisher, 1922; Noonan et al., 2006). H.J. Muller was the first to discover balancing selection from his study of balanced lethals in Drosophila (Muller, 1918). Adaptive variational maintenance by balancing selection may be achieved in a number of ways. In a mechanism known as over-dominance (also called heterozygote advantage), the individual who is heterozygous for a certain variant has a higher fitness (Fisher, 1922; Wallace, 1970). In another mechanism called negative frequency-dependent selection, rarer variants confer higher fitness. This leads to a fluctuation of a variant’s frequency in the population until an equilibrium is established, such that neither variant confers an advantage relative to the other (Smith Maynard et al., 1998; Takahashi & Kawata, 2013). Temporally varying selection, wherein the selection coefficient of an allele changes over time, can lead to the oscillation of this allele’s frequency over time (Abdul-Rahman et al., 2021; Wittmann et al., 2017). Spatially varying selection, wherein the selection coefficient of an allele varies across geography, may fix this allele locally in one niche and eliminate it in another, leading to the global persistence of variation at the locus (Hedrick, 2006; Levene, 1953; Saitou, Masuda, et al., 2021).

Unlike positive and negative selection, there is only a modest number of well-established instances of balancing selection (Charlesworth and Charlesworth 2016). In humans, these include polymorphisms of the *ABO* gene, which determine the A, B, and O blood groups (Ségurel et al. 2012), and polymorphisms in the major histocompatibility complex, which encodes cell-surface glycoproteins that display samples of peptides from within the cell on the cell’s surface (Takahata et al. 1992). Two variants of *ERAP2*, which too is involved in the antigen-presenting pathway, have been maintained under balancing selection as well (Andrés et al., 2010). The classic example of recent, shorter-term balancing selection in humans is the maintenance of sickle-cell alleles at the β-goblin locus (by over-dominance) in the regions of Africa where malaria is endemic (Allison, 1954a a, 1954b b; Hedrick, 2011). Similar reasoning applies to certain α-thalassemia alleles in parts of Southeast Asia where malaria is widespread (Qiu et al. 2013). In fact, the higher fitness of heterozygotes for thalassemia alleles in malaria-struck regions was presciently predicted by Haldane in 1949 (Lederberg, 1999). In the realm of structural variants, it was shown that a deletion spanning *LCE3B* and *LCE3C*, which is associated with psoriasis, has persisted in the human lineage for more than 700,000 years likely owing to balancing selection (Pajic et al., 2016).

So far, most systematic investigations into balancing selection in modern humans have focused primarily on genes (DeGiorgio et al., 2014) and on single nucleotide variants (SNVs), (Siewert & Voight, 2020); (Bitarello et al., 2018; Siewert & Voight, 2017). Additionally, some studies have focused exclusively on ‘long-term’ balancing selection wherein variants have been maintained in the human lineage since before the split from the chimpanzee clade (Leffler et al., 2013). Others have focused on short-term or population-specific balancing selection (Hedrick, 2011; Qiu et al., 2013). In order to identify potential targets of balancing selection that are structural in nature and that may not have been captured by such studies, we concentrate our efforts on autosomal deletion polymorphisms (>50bp) that have been maintained in the human lineage since before the split, approximately ∼700,000 years ago, of anatomically modern humans (AMHs) from the lineage that led to both Neanderthals and Denisovans (henceforth, collective referred to as archaic hominins). Targeting examples of such ‘medium-term’ balancing selection will likely allow us to capture more potential targets than could an exclusive study of either ‘long-term’ or ‘short-term’ balancing selection. Moreover, deletions are interesting in the context of selection. Since a given deletion affects more nucleotides than a single nucleotide variant (SNV), if a defined genomic sector is indeed of adaptive importance, deletions may have more profound functional consequences (Conrad et al., 2010; Saitou & Gokcumen, 2020). Such functional outcomes may translate into non-trivial selection coefficients either for or against the deletion. Additionally, deletions are relatively easier to genotype and analyze compared to other structural variants, which makes an evolutionary analysis involving deletions tractable.

## RESULTS AND DISCUSSION

### A significant portion of ancient polymorphisms may have evolved under medium-term balancing selection

Older polymorphisms may be more likely than newer ones to exhibit signatures associated with balancing selection because they have survived the stochastic fixation or elimination for extended periods. It is, therefore, possible that a certain proportion of human polymorphisms that are older than the AMH-archaic split (∼700,000 years) (**Figure 1A**) have been maintained by balancing selection. We tested this hypothesis by comparing the observed allele sharing between archaic hominins and AMHs to a neutrally expected distribution. If this proportion is significantly higher in the observed data than under the neutrally simulated data, and if we can reject other plausible explanations for the inflated sharing, we can conclude that some of the polymorphisms older than 700,000 years may have been maintained by balancing selection.

**Figure 1.**
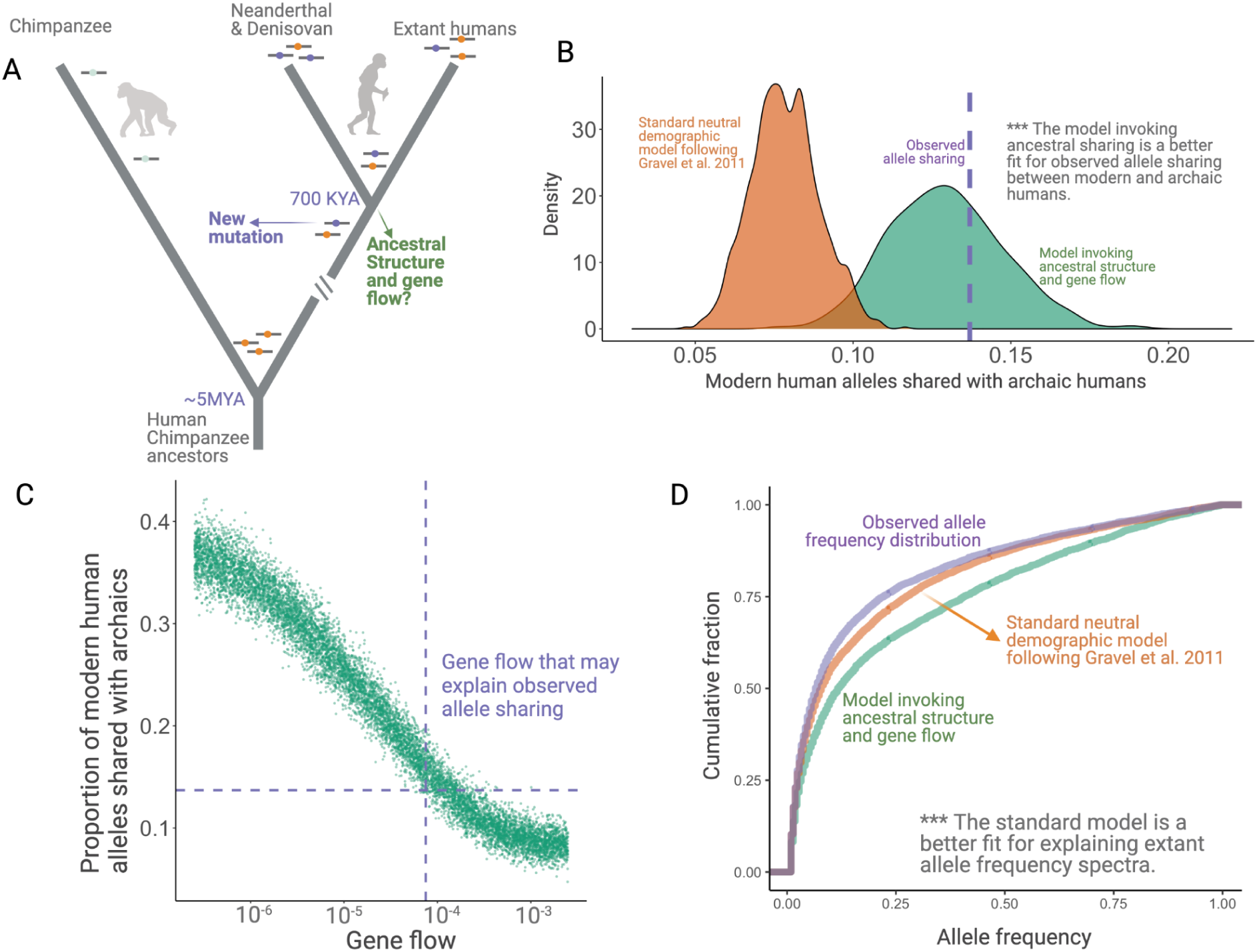
Excess of allele sharing between archaic hominins and modern humans. **A**. A schematic representation of derived “ancient” variations (purple) that emerged before the AMH-archaic divergence (and after hominin-chimp divergence) and have remained polymorphic in the modern human lineage. **B**. Allele sharing between modern humans and archaic hominins. The purple line is the observed allele sharing in Yoruba (YRI). The green and orange density plots indicate allele sharing in neutral simulations with and without ancestral structure, respectively. **C**. Results from simulations invoking ancestral structure with different gene flow levels (migrants per generation introduced into the populations) among the structures, indicated on the x-axis. **D**. Comparison of the allele-frequency spectra of simulated SNVs with observed SNVs. The purple, orange, and green lines represent allele frequency spectra in the YRI population using actual SNVs, neutral simulations without ancestral structure, and neutral simulations invoking ancestral structure, respectively.

For this test, we focused on 28,291 randomly chosen SNVs (minor allele-count > 1) in the Yoruba (YRI) population (1000 Genomes Project Consortium et al., 2015); see **Supplementary material** for a discussion of our rationale behind using SNVs rather than deletion polymorphisms, the variant class of our interest). We found that the derived SNVs shared with archaic hominins by common descent make up 13.7% (3,894 SNVs) of the total. To compare this number against the neutral expectations, we used *ms* (Hudson, 2002) to neutrally simulate 10,000 sets of 1,000 variants (see **Methods** for details). For these simulations, we used previously outlined neutral parameters determined for modern human populations by (Gravel et al., 2011). For the divergence times and effective population sizes of archaic hominins, we used parameters outlined elsewhere (Bergström et al., 2021; Mafessoni et al., 2020; Meyer et al., 2012; Prüfer et al., 2014). We found that the entire distribution of derived allele sharing with archaic hominins (median=7.8%) lies to the left of the observed sharing of 13.7% (**Figure 1B**). Therefore, we conclude that at least under this neutral model, the high proportion (13.7%) of polymorphisms that have been maintained for more than 700,000 years cannot be explained by drift alone.

Next, we asked whether we can reject plausible explanations of this excess of ancient polymorphisms other than balancing selection and the neutral model characterized by the parameters stipulated by Gravel et al. (2011). One alternative explanation comes from the evidence for introgression from early modern human ancestors into Neanderthals to the exclusion of Denisovans (Posth et al., 2017). Such admixture can increase the apparent allele sharing between Neanderthals and Yoruba. Nevertheless, we showed that there is still a marginal excess of SNVs (9.7%) wherein the derived allele is shared with the Denisovan genome relative to neutral expectation (**Figure S1**; p=0.052). We note that the allele sharing with this Denisovan individual may also be reduced due to a previously suggested superarchaic component in Denisova (Meyer et al., 2012; Rogers et al., 2020). Therefore, even if we concede that the overall excess sharing of SNVs in Yoruba with archaic hominins is due to introgression from early modern humans into Neanderthals, the excess of variants shared with Denisovans would remain partly unexplained.

The excess allele sharing mentioned above can also be caused if the modern and archaic lineages coalesce, on average, much farther into the past than in our simulations above. One way to achieve this is by invoking structure in the common ancestral population of archaic hominins and AMHs. This will effectively reduce the coalescence rate between the archaic and modern human lineages, thereby increasing their time to coalescence. To see what level of structure would be required to cause the observed excess, we conducted another set of simulations invoking structure in the population ancestral to AMHs and archaic hominins, while allowing gene flow between these latent subgroups (**Figure 1A**; see **Methods** for details). We found that the observed allele sharing between Yoruba and archaic hominins can indeed be explained by structuring the ancestral population into 3 distinct subpopulations, such that the fraction of each subgroup formed by the migrants of each of the other subgroups, every generation, is below 0.0075% (**Figure 1C**). However, the allele frequency spectrum for SNVs simulated with ancient population structure significantly deviates from the observed allele frequency spectrum in that the former overestimates the intermediate/common variants (**Figure 1D**). Therefore, invoking this kind of ancient structure to explain the excess allele sharing may be unrealistic.

This motivates our following analysis in which we test whether balancing selection may be acting on ancient (older than 700,000 years) polymorphisms and can explain, at least in part, the excess sharing of derived alleles with archaic hominins.

### Categorizing human deletions based on their evolutionary history

Having established that human polymorphisms older than ∼700,000 years likely include targets of ‘medium-term’ balancing selection, we next identified human deletion polymorphisms that have been maintained since before the AMH-archaic split. Since the vast majority of deletions in modern humans are derived relative to chimpanzees (**Supplementary material**), this could be accomplished by identifying AMH deletions that are shared with archaics by common descent. To this end, we used the 1000 Genomes Project Phase-3 dataset (1000 Genomes Project Consortium et al., 2015), filtering it to focus on 4,863 human deletion polymorphisms that are in linkage disequilibrium (LD, r^2^ > 0.9) with at least one SNV (**Table S1**). We imposed this linkage requirement because linked SNVs enabled us to distinguish the shared deletions that are introgressed or recurrent from those that are identical-by-descent. For this analysis, we used deletions that were present, with an allele-count greater than 1, in the YRI (Yoruba), CEU (Utah residents with Northern and Western European ancestry), and CHB (Han Chinese in Beijing) combined. None of these deletions overlap with polymorphic inversions in humans reported in the 1000G Phase-3 dataset.

We found that 575 (11.8%) AMH deletions in the *deletion dataset* were shared with archaic hominins (**Figure 2**). Specifically, we genotyped all the deletions in our dataset in the four available high-coverage (∼30X) archaic hominin genomes (Mafessoni et al., 2020; Meyer et al., 2012; Prüfer et al., 2014, 2017), using a read-depth based pipeline (**Table S2**). We identified 53 instances of independent emergence (*recurrent deletions*) in modern humans and archaics, wherein no SNV in LD with the deletion in modern humans accompanied the deletion in the archaic genomes. In parallel, we identified 92 deletions *introgressed* from archaic hominins into modern humans using the linked SNVs that were present in previously identified introgressed haplotypes (R. O. Taskent et al., 2017; Vernot & Akey, 2014). By this process of elimination, we found that 430 (8.8% of the total) shared deletions are ancient, *i.e*., they are shared with the archaic lineage by common descent and thus emerged at least ∼700,000 years ago.

**Figure 2.**
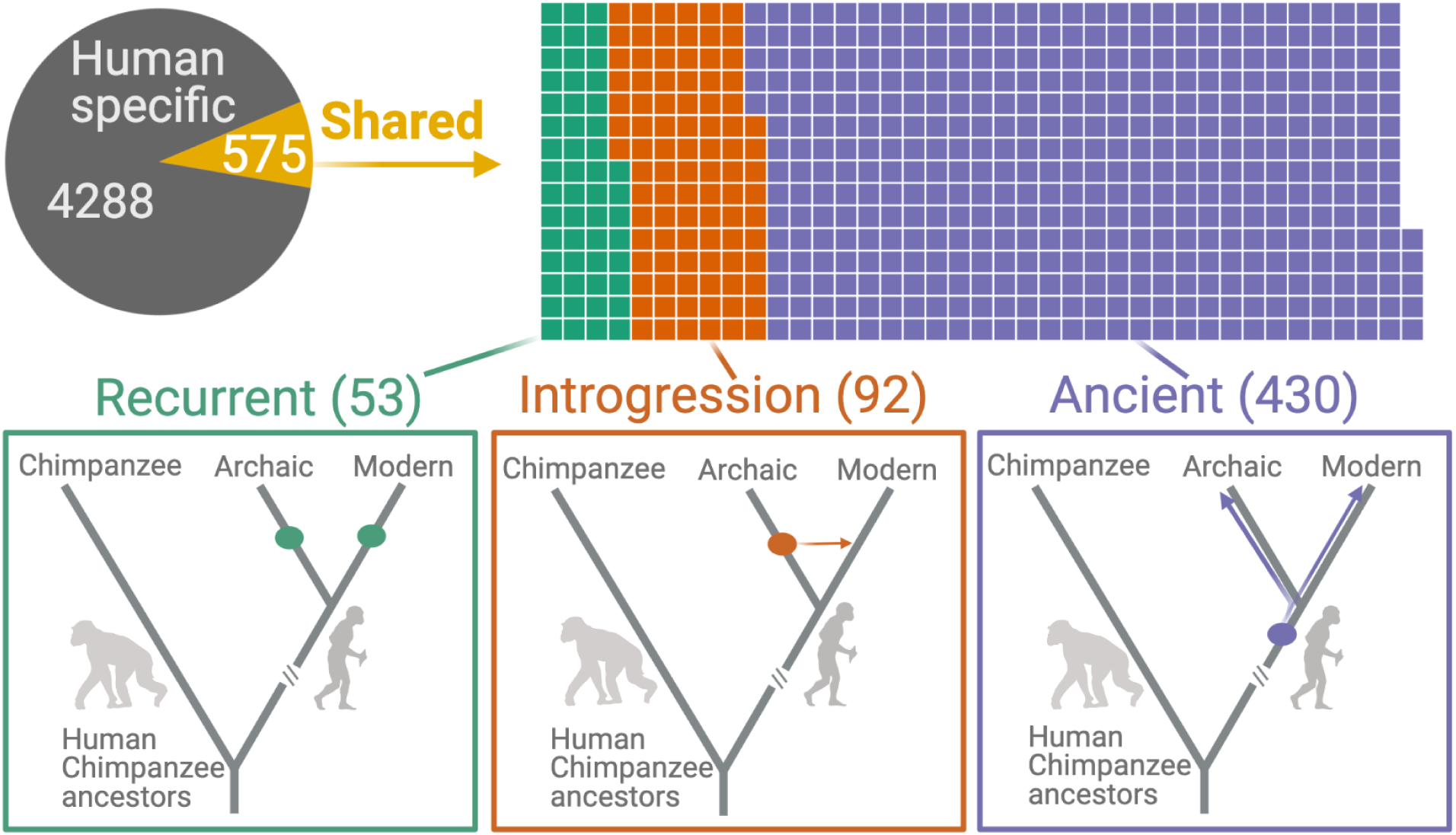
Deletions in modern humans that are shared with archaics. The top panel shows the categorization of deletion polymorphisms as human-specific, recurrent, introgressed, versus ancient. The evolutionary histories of shared deletions are summarized schematically at the bottom panel.

To confirm that our pipeline for identifying ancient deletions has high accuracy, we calculated the ages for all deletions. If our genotyping pipeline and categorization of shared deletions (as recurrent, introgressed, versus ancient) is sound, we should expect a specific pattern of average ages across the categories of deletions. In particular, we expect that ***Age****(human-specific)* ≈***Age****(recurrent) <* ***Age****(introgressed) <* ***Age****(Ancient)*. We estimated the ages of the deletions in the *deletion dataset* using two methods: 1) Human Genome Dating (Albers & McVean, 2020); and 2) Relate (Speidel et al., 2019) **(see Methods)**. Both methods yielded the expected pattern of ages across the categories of deletions **(Figure 3, Figure S2)**. We found that the median age for the ancient candidate deletions, using Relate, is 1,004K years. About 15% (63) of these deletions are older than 2 million years. In contrast, the median age of deletions that are not shared with archaics by common descent is ∼239K years. As such, we infer that our genotyping pipeline and deletion categorization approach are sound.

**Figure 3.**
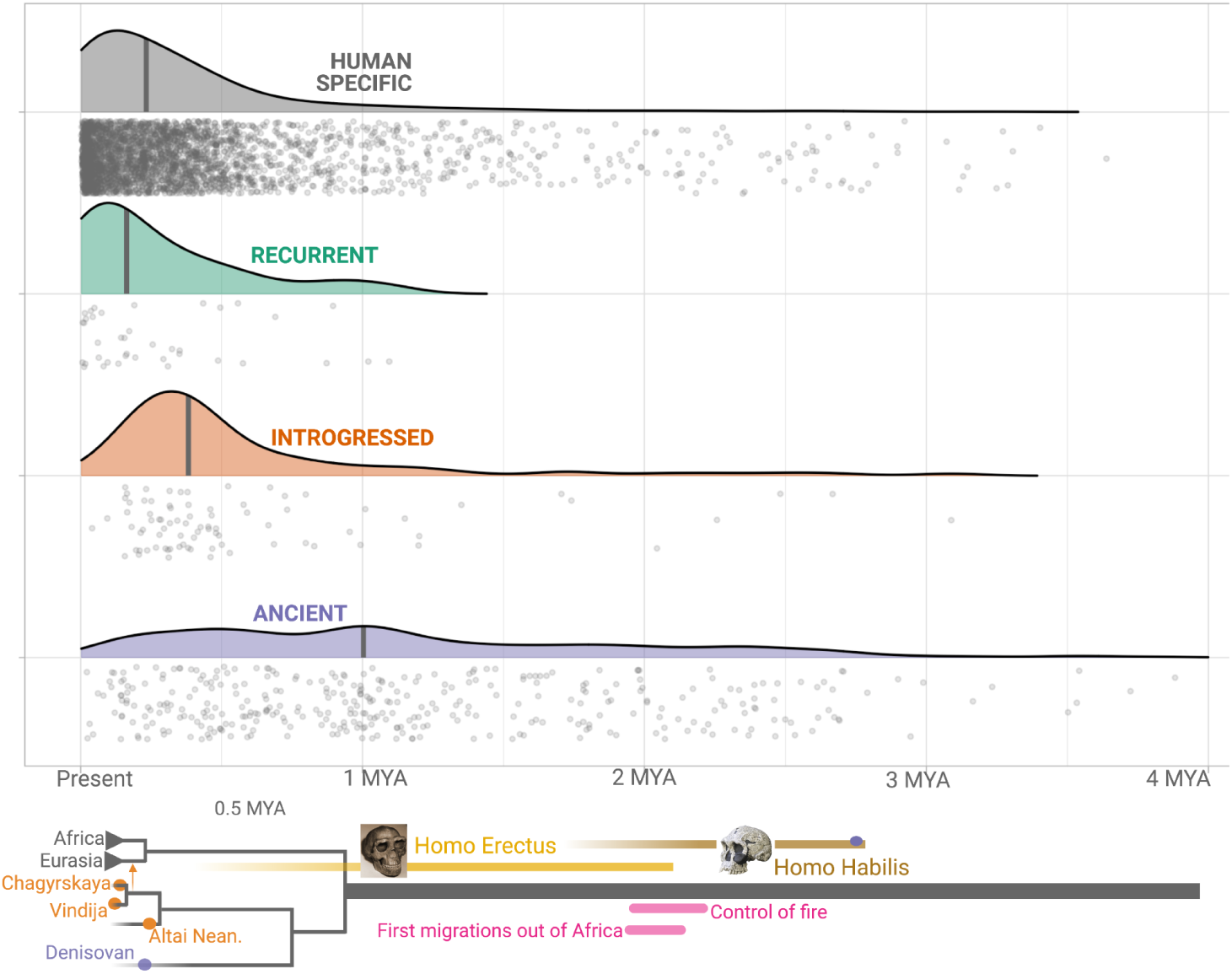
Age estimates of the haplotypes harboring polymorphic deletions. The x-axis shows the age estimates of haplotypes harboring the deletions, obtained using Relate. For orienting the reader regarding the age of these variants, we provide below a schematic phylogeny representing recent human evolution.

### Ancient deletion polymorphisms are more likely to be targets of balancing selection than younger ones

We empirically tested our hypothesis that ancient deletion polymorphisms are more likely than younger ones to have evolved under balancing selection using the std***β***2-statistic. A recent and robust measure of balancing selection (Siewert & Voight, 2017), std***β***2, measures the weighted average of the number of flanking derived variants, where weights are the similarity in frequency between the core allele and the flanking variants (**Figure 4A**). Suppose a variant has evolved under balancing selection. In that case, we expect an excess of variants surrounding the core variant that are at the same frequency as the core variant itself. If a deletion emerges and the resulting polymorphism is subject to balancing selection, the deletion will rise in frequency until it reaches a certain equilibrium frequency. New SNVs will emerge on the haplotypes carrying the deletion. Some of these SNVs will drift upward in frequency, but since these SNVs will be linked to the deletion, they too can only rise to the equilibrium frequency of the balanced deletion (Siewert & Voight, 2017, 2020). We refer to this type of drift as *Goldilocks drift*, since the linked SNVs drift upward to the “just-right” equilibrium frequency of the balanced deletion. Goldilocks drift thus leads to allelic class build-up, which refers to a situation involving the fixation of many flanking variants within the set of haplotypes carrying the deletion. Therefore, a high std***β***2 value for a variant implies that it is close to a target of balancing selection. We observed that std***β***2 estimates for ancient deletions are significantly larger than other deletions across YRI, CEU, and CHB populations (p<10^−7^, Wilcoxon) (**Figure 4B**). These empirical results provide evidence that ancient deletions are enriched for targets of balancing selection. Our results are consistent with other recent studies (Soni et al., 2021) that have argued that the role of balancing selection in explaining the maintenance of common variation in the human lineage is underappreciated.

**Figure 4.**
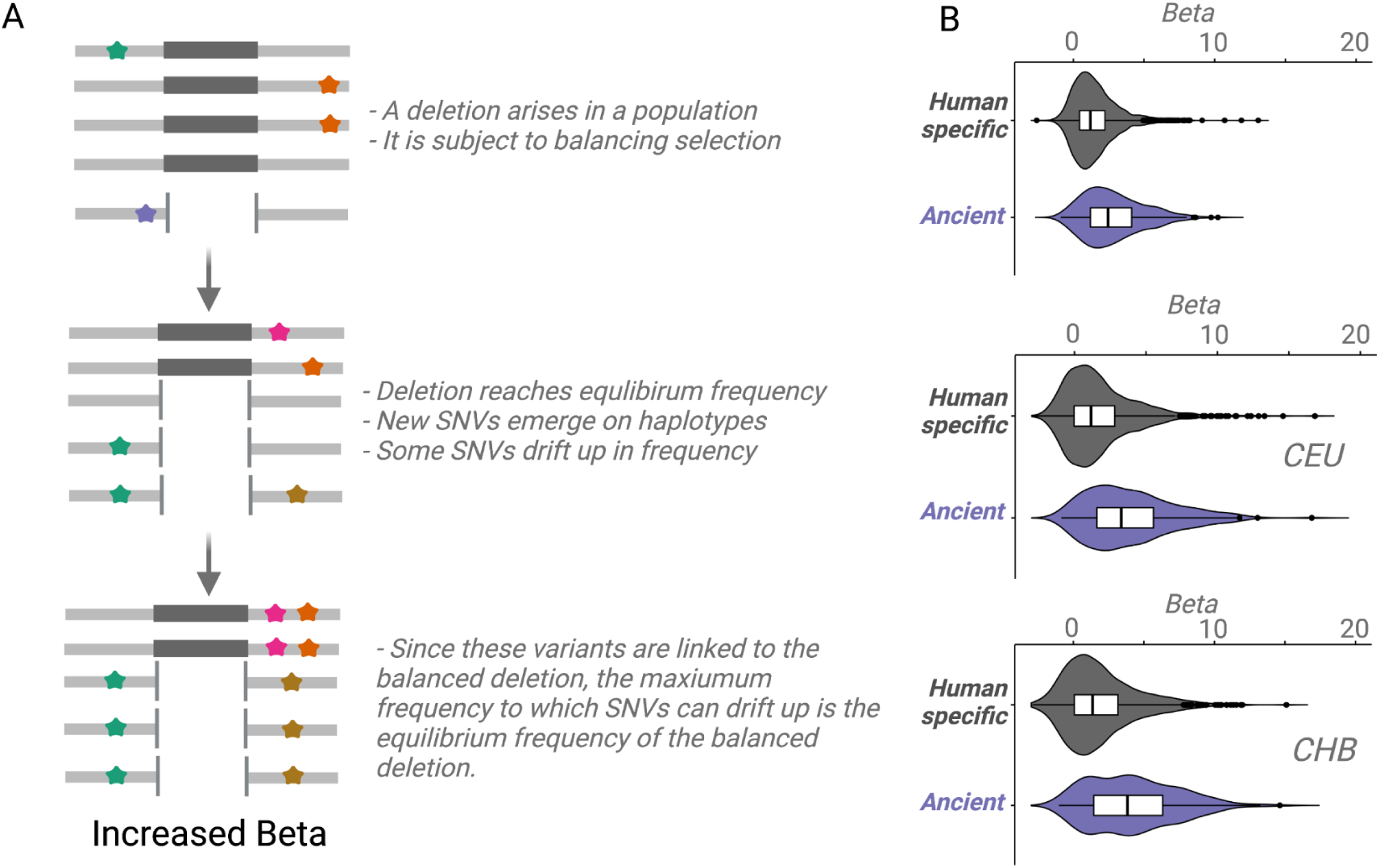
An empirical assessment of putative balancing selection among ancient deletions. **A**) The conceptual framework in which ***β*** statistic works. The last step demonstrates ‘Goldilocks’ drift (the process that results in the allelic class build-up). **B**) A box plot for beta values calculated for SNVs in linkage disequilibrium with AMH-specific, ancient polymorphic, versus ancient fixed deletions. Higher std***β***2 values for older deletions represented in purple empirically show that older deletions are significantly enriched for targets of balancing selection. All comparisons are significant, p<10^−7^ (Wilcoxon).

### Strong overdominance is rare among deletion polymorphisms

Having established that ancient deletion polymorphisms appear enriched for targets of balancing selection, we wanted to investigate whether classical overdominance is a common mechanism underlying this observation. To accomplish this, we first identified the genomic signatures that we expect to see in a region where a polymorphism has evolved under overdominance, and then we look for these signatures among ancient deletions. To identify the signatures of overdominance, we simulated sequence evolution under neutrality and overdominance, using a variety of selection coeffiecients, for variants that emerged one million years ago. We asked whether we can distinguish between neutrality and overdominance by calculating several population genetic statistics on sequences generated from the neutral and overdominance simulations, including Tajima’s D, π, θ, etc. (see **Methods** for full list). We found that none of these statistics alone can distinguish overdominance from neutrality, even for strong selection coefficients. We further found that even when combined, using principal component analysis, these statistics could not distinguish neutrality from balancing selection (**Figure 5A**).

**Figure 5.**
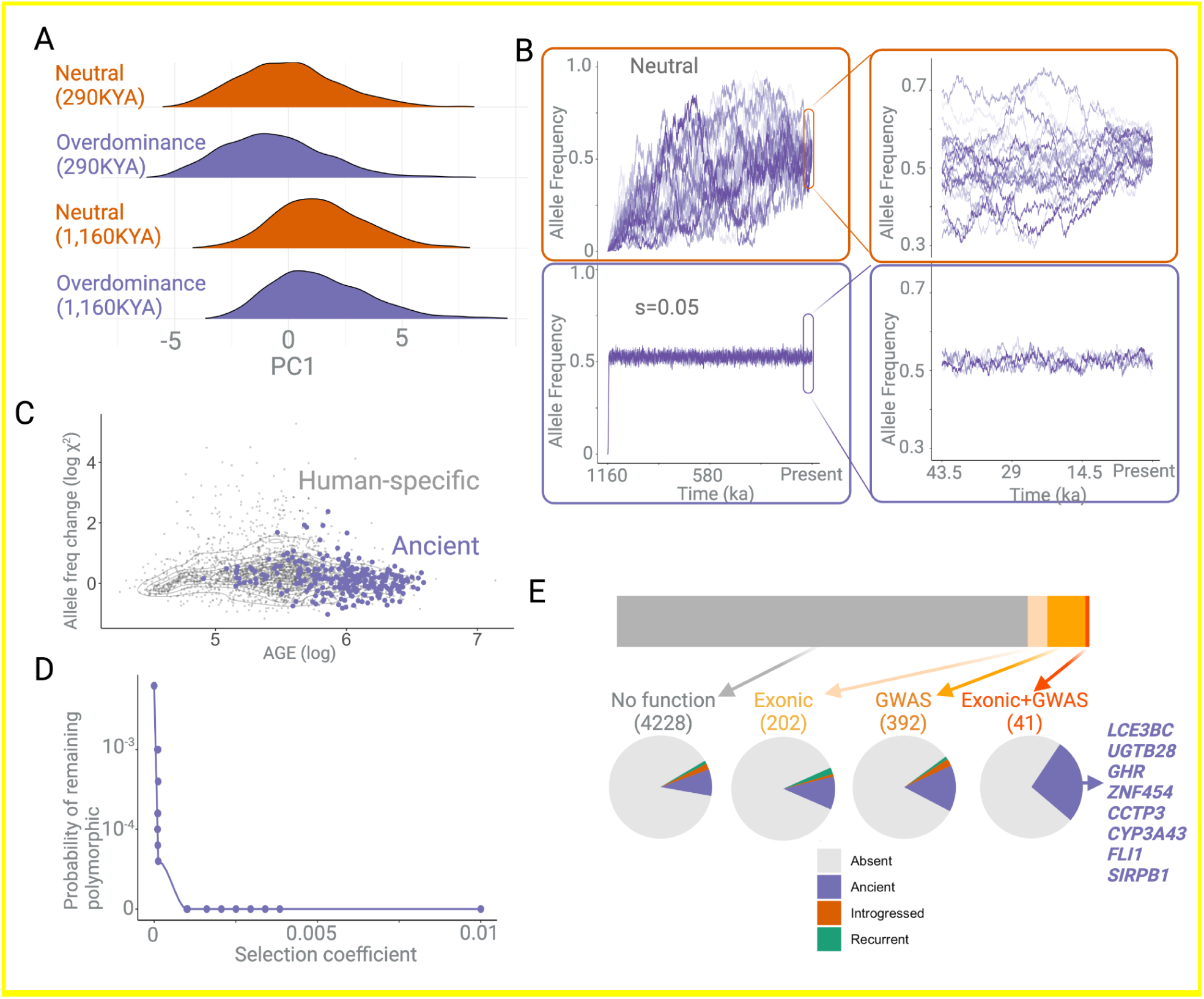
**A**. First principle component of multiple summary statistics based on simulations ran on neutral versus overdominance (s=0.05) expectations. There is no discernible difference between overdominance and neutrality within the time-frame of these simulations. **B**. The allele frequency trajectories of variations over 1,000,000 years evolving neutrally (top), versus under overdominance (bottom). The x-axis represents the time since the emergence of a variant in years, assuming a 29 year generation time. The right panel is a zoomed-in version of the same allele frequency trajectories in the last ∼50 thousand years. **C**. Estimated measure of allele frequency change (χ^2^) between 50,000 and 5,000 years before present as a function of allele age in years (x-axis). The allele frequencies of ancient deletions are not necessarily more stable in this time frame than genome-wide expectations. **D**. The expected number of alleles that remain in the populations over the last 1,000,000 years under different negative selection pressures. **E**. The proportion of deletions that are “functional” (i.e., exonic or associated with GWAS traits).

The one marked distinguishing factor between neutral and balancing selection scenarios that the simulations highlighted is that the allele frequency trajectories are quite different for the two cases (**Figure 5B**). Under neutrality, as expected, a random change in allele frequency in every generation generated elevated noise across time in allele frequency trajectories. In contrast, under overdominance, the allele frequency rapidly increases (similar to a selective sweep) until it reaches an equilibrium frequency, whereafter it remains remarkably stable across time. To ascertain if overdominance may be a common mechanism of evolution among ancient deletions, we inferred the allele frequency trajectories of ancient deletions using *Relate* (Speidel et al., 2019) and quantified the allele frequency change between 5,000 and 50,000 years ago by squaring the standardized allele frequency difference (χ^2^) (**Methods**). If indeed, the maintenance of older alleles is partly due to overdominance, we expected such deletions to have more stable allele frequencies across time, leading to a smaller χ^2^ value. However, we found no general trends that suggest that any of the ancient deletions show significantly lower allele frequency changes over time, as compared to genome-wide expectations (**Figure 5C**). Consequently, at least with our current resolution of allele frequency trajectory estimation, we found no evidence for a strong overdominance effect among ancient deletion polymorphisms in modern human populations. However, this finding depends on the specific parameters that we assumed in our simulations and the power of our analysis.

### Functional ancient deletions are likely targets of balancing selection

Based on previous work, we expect that deletions are more likely than SNVs to be under negative selection (Conrad et al., 2006; Kondrashov, 2017; Lin et al., 2015; Lin & Gokcumen, 2019). To investigate the magnitude of this effect, we compared the proportion of SNVs and deletion polymorphisms that are ancient. Applying the same bioinformatic pipeline to identify ancient polymorphisms in both cases, we found that 13.7% of SNVs and 8.8% of deletion polymorphisms are ancient. This result alone suggests that deletion polymorphisms are more likely than SNVs to be eliminated by negative selection, a trend that we expect to be more pronounced with increasing ages of polymorphisms. The greater intensity of negative selection acting on deletions implies that deletions are, in general, more deleterious than SNVs. It follows that non-ancient deletions currently segregating in human populations are more likely to be deleterious than SNVs (Kondroshov, 2017), and therefore, more likely to be in the process of being eliminated by negative selection.

Next, we conducted simulations to determine how different magnitudes of negative selection acting on variants that emerged 1 million years ago affect the proportion of variants that are eliminated. Our simulations suggest that even relatively weak (s = 10^−4^) negative selection acting on variants that emerged 1,000,000 years ago should eliminate more than 99.9% of polymorphisms **(Figure 5D; see methods for details)**. This suggests that old polymorphisms only rarely survive if they evolved under negative (or by proxy positive) selection. Therefore, we hypothesize that most surviving ancient deletion polymorphisms have either evolved under neutrality or some sort of balancing selection. It follows that the small proportion of ancient deletion polymorphisms that are functional, and therefore, are likely to have fitness effects, are prime candidates for being targets of balancing selection.

Given that we have not found evidence for strong overdominance among ancient AMH deletion polymorphisms, we conclude that the vast majority of the instances of such medium-term balancing selection likely involve temporally and spatially variable selection. This is consistent with our locus-specific analyses of ancient deletion polymorphisms. For example, we recently reported that the deletion of the third exon of the human growth hormone receptor gene (*GHR*) has evolved under temporally and geographically variable adaptive constraints (Saitou,

Resendez, et al., 2021). These ancient deletions are common and old, and also exhibit high population differentiation. Collectively, we argue that such adaptive maintenance of ancient, functional alleles may be due to varying selection trends across geographies and time.

Selection should only act on a region of the genome by means of the phenotypic function associated with that region. It follows then that any adaptively maintained ancient polymorphisms must be functional. To this end, we ascribed functions to deletions by identifying SNVs linked to deletions that are significantly (p<10^-8) associated with human traits (http://www.nealelab.is/uk-biobank/; see **Table S3** for a consolidated list of all statistically significant association). We found 63 such ancient deletion polymorphisms, including the Crohn’s disease-associated deletion upstream of the *IRGM* gene (McCarroll et al., 2008). We also identified the deletions that intersect with protein-coding sequences. We found that, among all the 4,863 deletions that we investigated, 243 (5%) are exonic, 433 (8.9%) are in haplotypes that harbored GWAS hits, and 41 deletions (0.84%) are both exonic and present in haplotypes with GWAS hits. To be conservative, we only considered a deletion functional if it was both exonic and associated with a phenotype. Of all the 41 polymorphic functional deletions, 11 (27%) were ancient (**Figure 5E**). These include well-known polymorphisms, such as the psoriasis-associated deletion of the *LCE3B* and *LCE3C* genes, which was previously argued to be evolving under balancing selection (Pajic et al., 2016). These results suggest that at least 27% (the proportion that is ancient) of common functional deletion polymorphisms may have been evolving under balancing selection.

### Functional ancient deletions are related to immune and metabolic function through varied mechanisms

To understand the functional effects of ancient deletions, we investigated the UK BioBank GWAS traits associated with them (**Figure 6a**). We found that ancient deletions primarily affect blood, immunity, and metabolism traits. Previous studies also highlight these functional categories in shaping recent human evolution (Andrés et al., 2009; Bitarello et al., 2018; Fumagalli et al., 2009; Leffler et al., 2013). A focused literature review and analysis of functional effects associated with the haplotypes harboring these deletions reveal multiple mechanisms through which they affect function. We highlight some of these insights below.

**Figure 6.**
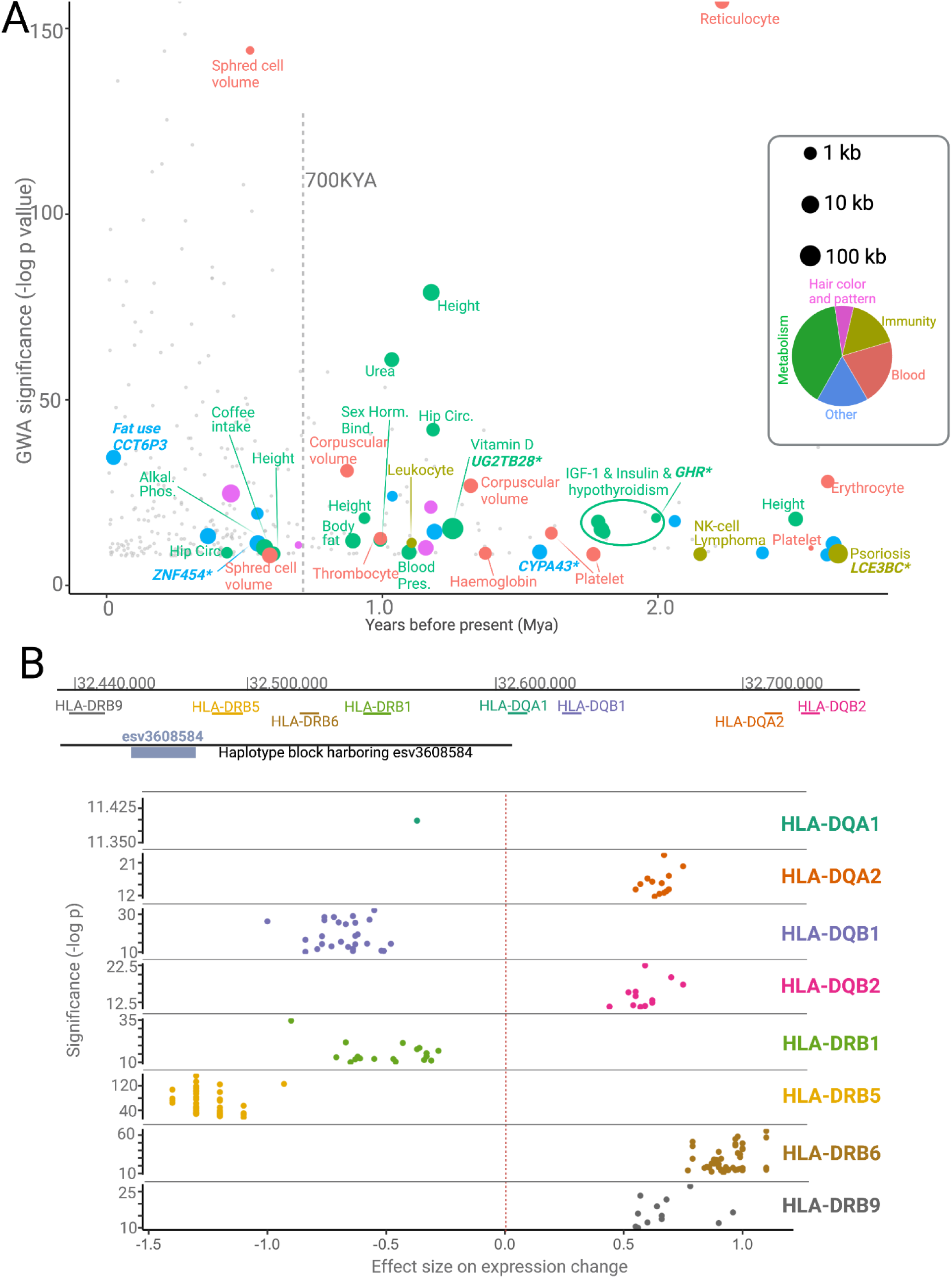
**A**. The significance of association for the deletions that reside in haplotypes associated with GWAS traits as a function of their emergence time. Gray points indicate human-specific deletions, where other colors indicate ancient deletions categorized functionally. **B**. The size and significance of the effect of the haplotype harboring esv3608584 on the expression of nearby *HLA* genes.

Firstly, whole gene deletions may affect the function of entire environment-interacting gene families. We found two ancient whole gene deletions: esv3587563 (deleting *LCE3B* and *LCE3C*), and esv3600896 (deleting *UGT2B28*). The members of both the *LCE3* and *UGT2B* gene families have been shown to likely be evolving under adaptive forces, and mediating immune-response and steroid metabolism, respectively (de Guzman Strong et al., 2010; Pajic et al., 2016; Starr et al., 2021; Xue et al., 2008). The functional consequence of whole gene deletions is, of course, loss of function of the deleted genes. In addition, esv3587563 is associated with an increase in the expression of LCE3A *(de Guzman Strong et al*., *2010; Pajic et al*., *2016)*, while esv3600896 is associated with an increase in the expression of *UGT2B11*. Thus, we propose that whole-gene deletions of members of environment-interacting gene families may lead to the functional “fine-tuning” of the entire gene family.

Secondly, we revealed dozens of potentially adaptive ancient deletions that mediate gene regulation and are associated with human traits. For example, we found multiple ancient deletions that are proximal to the *HLA* locus, mostly associated with immune-related phenotypes, and affect the expression levels of various *HLA* genes (**Figure 6B**). More surprisingly, we observed that the same ancient deletions also lead to the expression of different isoforms of HLA genes. Using the GTeX database, we found at least four other instances where ancient deletions lead to the expression of different isoforms, including deletions affecting the *HLA-DRB1-6, HLA-DOB, SIRPB1, GHR* and *CYP3A43* genes. We recently showed that the ancient deletion of the third exon of the growth hormone receptor gene leads to the expression of a smaller version of growth hormone, which may be adaptive in times of starvation (Saitou, Resendez, et al., 2021). The *SIRPB1* gene encodes a glycosylated transmembrane receptor protein (Kharitonenkov et al., 1997) and its different isoforms may lead to the recognition of different pathogens. Similarly, *CYP3A43*, a member of the cytochrome p450 gene family, is involved in metabolizing external substances, and genetically determined isoforms contribute to its functional variation in humans (Agarwal et al., 2008). Thus, ancient deletions that lead to specific isoform expression may have been adaptively evolving to adjust the function of environment-interacting genes across both geography and time. It is important to acknowledge that these non-exonic deletions may not be the causal variant in the associated haplotypes. Nevertheless, the full extent of deletion polymorphisms shaping the expression levels and sculpting the isoform diversity at the genetic level remains a fascinating area of future research.

## CONCLUSION

This manuscript asked whether adaptive forces have maintained ancient deletion polymorphisms in humans. We provide evidence that there is a greater number of ancient polymorphisms in modern human genomes that predate the divergence of humans from Neanderthals and Denisovans, relative to the neutral expectation. We identified 63 ancient deletions associated with known human traits. Using simulations and empirical data, we provide evidence for the notion that balancing selection is likely a considerable force shaping the extant functional deletion polymorphisms. Specifically, we show that about 27% of such functionally relevant deletions are ancient. Our results suggest that classical overdominance may not explain the adaptive forces acting on these ancient deletions. Instead, geographically and temporally variable, and frequency-dependent selection may underlie the maintenance of ancient functional deletions. Mechanistically, in addition to previously defined functional effects such as whole gene deletions and regulatory variants, we highlight multiple instances where ancient deletion polymorphisms lead to the expression of different isoforms. Overall, our study contributes to the growing evidence that balancing selection may be a considerable force in shaping common, functional deletion variation in extant human populations.

## DATA AVAILABILITY

All data generated can be found in the supplementary files. The codes that we used to generate our datasets and simulations can be found either in Methods or in our GitHub page **(https://github.com/GokcumenLab)**.

## SUPPLEMENTARY TABLES

**Supplementary Tables** can be found in this link - https://www.dropbox.com/s/nwpdqo5lse51c6u/supplement.zip?dl=0

## ACKNOWLEDGEMENTS

We thank Dr. Victor Albert and Dr. Vincent Lynch for their careful reading of this manuscript. We acknowledge Petar Pajic and Charikleia Karageorgiou for their insightful discussions throughout the development of this project.

## ETHICS

All data used in this study has been previously published in peer-reviewed journals.

## METHODS

### Identifying deletions in archaic genomes

The identification of deletions in archaic hominins was predicated on the concept that a deletion in an archaic hominin would correspond to a low read depth in the window of deletion in the hominin’s genome. We started with two main types of input files: 1) The VCF file for the 1000 Genomes Phase 3 dataset; and 2) A BAM file for each high-coverage archaic hominin.

The 1000G phase-3 VCF file was obtained from https://www.internationalgenome.org/data. This includes 84.4 million variants from 2504 individuals across 26 populations. This VCF file was filtered to retain only biallelic autosomal deletions that were present in the CHB, YRI, or CEU populations. The file was then converted to a BED file (with three tab-separated columns representing chromosome numbers, start positions, and end positions of deletions) using a script written in BASH. This BED file contained the information about the start and end coordinates of the deletion. This amounted to 32,154 deletions. Note that for the analysis described in the main text, only 7,378 of these deletions (those that had an allele-count > 1 in YRI, CEU, and CHB combined) were used.

The sequence files for archaic genomes mapped to hg19 were obtained from https://www.eva.mpg.de/genetics/genome-projects.html?Fsize=0%2C%252%27A%3D0. These BAM files containing mapping information (such as the start and end coordinates of the part of the genome to which a read maps) were converted to BED files. This was done using the `bamToBed` command in the `bedtools` module (Quinlan & Hall, 2010). We then used the two types of BED files to count the number of reads for each archaic genome that mapped to a region of the genome that is polymorphically deleted in modern humans. In order to achieve this, we used the `intersectBed` command with the `-c` option within the `bedtools` module. This command counts the number of reads in an archaic genome that intersects with the region of the genome harboring a deletion polymorphism in modern humans.

Next, for every archaic genome, we normalized by size the number of reads at each window of deletion using the following formula

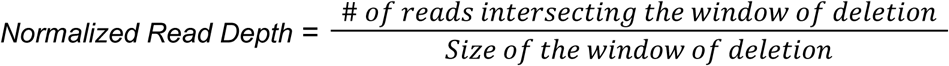

For each archaic genome, we wanted to calculate the Z-scores of the normalized read depths across all windows of deletion, and classify a window as a deletion if the normalized read depth was below a certain threshold. To prevent outliers from affecting measures of central tendency and spread, and therefore the Z-score threshold, we use the modified Z-score to classify a region as deleted or non-deleted in an archaic genome. The modified Z-score uses median (as opposed to mean) and median absolute deviation (as opposed to standard deviation) to calculate the Z-score. For a given archaic genome, the modified Z-score of the normalized read depth at the i^th^ window of deletion is given by

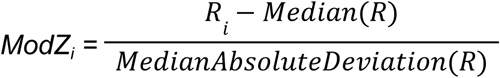

where R_i_ denotes the normalized read depth at a given window of deletion, and R denotes the random variable representing normalized read depth.

(Iglewicz & Hoaglin, 1993) have suggested that a threshold ±3.5 is reasonable for outlier detection using modified z-scores. Nevertheless, for our purposes, we used a more convervative threshold of -5, which we deemed more-appropriate based on spot-checking. For example, if the modified Z-score (of normalized read-depth) at a window of deletion was less than -5 in the Vindija Neanderthal, that window was classified as deleted in the Vindija Neanderthal. The distributions of these modified Z-scores across windows of deletions for the four high-coverage archaic genomes are illustrated in figure **S3 a-d**. All calculations downstream of obtaining the raw numbers of reads from archaic genomes intersecting with the windows of deletion, were performed using a script that we wrote in R. The read-depth analysis was done using all 32,154 AHM deletions (results for the status of these deletions in the four high-coverage archaic genomes are available in **Table S2**).

### Identifying SNVs that are in linkage disequilibrium with deletion in modern humans

We subsetted the 1000G phase-3 VCF files (there is a separate file for each chromosome) obtained from https://www.internationalgenome.org/data to retain individuals only from the CEU, CHB and YRI. This filter was applied using the `--keep` option in the module VCFtools (Danecek et al., 2011). All variants that had a minor allele count of less than 2 were eliminated using the `--mac` filter in VCFtools. We used the resulting VCF files to identify SNVs linked to each of the 7,378 autosomal biallelic deletions (this list had been created earlier as described in a previous section) with allele-count > 1 within a 50kb radius of the deletion. We did this using the --hap-r2-positions and --min-r2 0.9 flags in VCFtools. For each autosomal biallelic deletion with allele-count > 1, this gave us a list of SNVs linked to the deletion with r^2^ > 0.9, if such SNVs existed. At least one such linked SNV existed for 4,863 deletions. We called this set of deletion the ‘deletion dataset’ and based all our downstream analysis on it.

It is important to describe why we only focused on deletions with identifiable linked variants (most of them SNVs) flanking the deletions. We can only eliminate potentially introgressed deletions by checking whether at least one of the SNVs linked to the deletion is already known to be introgressed. Moreover, we can confirm whether a deletion shared between archaics and moderns is identical by descent (thereby eliminating recurrence) by checking whether the same SNVs accompany the deletion in moderns and archaics. This filtering would not be possible if our deletions were not flanked by variants linked to them. We have no reason to believe that eliminating deletions with no linked SNVs introduces a systematic bias in inflating or deflating the proportion of human polymorphic deletions that are shared (by common descent) with archaics. However, a shortcoming of this approach is that it fails to capture any balanced deletion that is not in linkage disequilibrium with at least one SNV.

### Eliminating instances of recurrence and introgression

Our method for identifying deletions in archaic hominins yielded 619 human polymorphic deletions that are also present in at least one archaic hominin genome. In order to ensure that we do further analysis only on deletions that are shared with archaics by common descent, we wanted to eliminate shared deletions that were recurrent or introgressed.

We removed recurrent deletions (those emerging in moderns and archaics independently) by retaining only those shared deletions for which at least one allele linked to the deletion in modern humans was also present in at least one archaic genome that harbored the deletion. To do this, we needed to know whether each linked variant is present or absent in the archaic genomes. We started with two types of inputs: 1) VCF files for each of the archaic genomes (mapped to hg19) and 2) a file containing all the variants (SNVs) linked to human polymorphic deletions. We filtered the VCF files to include only the SNVs linked to shared deletions. This was done using the `--positions` flag in VCFtools. The presence or absence of every linked variant was then determined using the vcf files for the archaics. The procedure was implemented using an AWK script. 76 shared deletions were classified as ‘recurrent’ using this approach.

In order to eliminate introgressed shared deletions, we used the results published by (O. Taskent et al., 2020). In their study, the authors had identified introgressed haplotypes in Eurasians using the S* statistic. They had also published a list of S*-significant SNVs that characterize introgressed haplotypes. We stamped out the shared deletions that were absent in Yoruba and for which at least one linked allele was among the S*-significant variant described in the study mentioned above. We thus eliminated 93 deletions that were likely introgressed from archaic hominins into modern humans.

### Ascribing phenotypic relevance to deletion

In the main text, we claimed that human polymorphic deletions that are shared with archaic hominins and have functional relevance will have an elevated probability of being targets of balancing selection.. This is the case because balancing selection can only affect a locus by acting on the phenotypes associated with the locus. Therefore in order to get a list of prime candidates, we set out to ascribe phenotypic effects to each of the 6,635 deletions in the deletion dataset. We used two criteria to ascribe phenotypic relevance to deletions: 1) any deletion intersecting with an exon was classified as phenotypically relevant; and 2) any deletion for which at least one linked variant was associated with a phenotype was classified as phenotypically relevant. All other deletions were classified and phenotypically insignificant.

In order to identify deletions that intersect with exons, we started with the genome annotation file download from https://hgdownload-test.gi.ucsc.edu/goldenPath/hg19/bigZips/genes/hg19.refGene.gtf.gz. Using an AWK script, this GTF file was then converted to a BED file containing five columns: 1) annotation’s chromosome number; 2) annotation’s start position; 3) annotation’s end position; 4) gene name; and 5) type of feature. Only the rows wherein the type of feature was ‘exon’ and the chromosome number was between 1 and 22 were retained. All repeated entries (rows) were eliminated. The resulting file contained only columns 1) to column 4). We then used a BED file containing information about the human polymorphic deletions in our deletion dataset and the BED file mentioned above to identify deletions spanning exons. This was done using the intersectBed option with -wa and -wb flags in the BEDtools module. On the resulting file, we used the groupby tool with the ‘-o freqdesc’ flag in the BEDtools module in order to obtain a file containing the names of the genes (and the number of exons within each intersecting gene) that overlap the deletions. 404 (6.1%) of the 6,635 deletions in the deletion dataset were exonic.

The second method to ascribe phenotypic relevance to deletions was to use results from previously published Genome Wide Association Studies (GWAS). We used a publicly available catalog of GWAS results based on the UK BioBank data (http://www.nealelab.is/uk-biobank/). In particular, we used data for 4,113 traits. For each trait, we used data that produced results using both sexes. For continuous traits, we used the raw version of the data, as opposed to the inverse rank normalized version. For each trait, only those SNVs were retained that were associated with the phenotype with a p-value less than 10^−8^. Then, for each of the 6635 deletions in our deletion dataset, we checked if any of the linked SNVs were among the SNVs that were significantly (p<10^−8^) associated with a phenotype. We then obtained the phenotype that was linked to one of the linked SNVs with the lowest p-value, and ascribed it to the deletion. Thus, 484 (7.3%) of the 6,635 deletions in the deletion dataset had phenotypic associations.

### Proportion of allele sharing in neutral simulations versus observation

The purpose of simulations was to establish whether there is an excess of polymorphisms maintained in the human lineage for more than 700,000 years, relative to the neutral model. If a derived allele is shared between humans and archaics by common descent, it emerged prior to 700,000 years ago. We compare the proportion of polymorphisms older than 700,000 years (derived allele shared with archaics) under the neutral model (represented by neutrally simulated SNVs) against the proportion observed in real-life (represented by randomly chosen Yoruba SNVs). If this proportion is significantly higher in real-life than under neutrality, we will have shown that choosing polymorphisms that have been maintained for more than 700,000 years potentially enriches for the target of balancing selection.

We obtained SNVs from the 1000G phase-3 vcf files (1000 Genomes Project Consortium et al., 2015) for the analysis described above. Using a script written in AWK (Aho et al., 1978), we subsetted the vcf files to retain only those biallelic SNVs that contained information about the ancestral/derived status of the two alleles. We used the --keep option in vcftools (Danecek et al., 2011) to retain individuals only from the Yoruba population. We then used the --mac option to retain only those SNVs for which the minor allele count was greater than 1. The allele-count filter was used to exclude singletons which could create spurious results for the linkage disequilibrium analysis described below. On the resulting vcf files, we used the SelectVariants tool with the --select-random-fraction option in GATK (Van der Auwera & O’Connor, 2020) to randomly retain 0.25% of the variants. This resulted in a set of 38,231 SNVs. Next, we investigated whether the random SNVs are in linkage disequilibrium with any other SNVs in their vicinity. In particular, we used the --hap-r2-positions and --min-r2 option in vcftools, along with the ancestral/derived status of alleles, to retain only those random SNVs wherein the derived allele was in linkage disequilibrium (r^2>0.9) with the derived allele of another SNV within a 50kb radius of the random SNV. This resulted in 28,491 SNVs. We only retained the random SNVs with linked variants in their vicinity to rule out cases of recurrence of the SNV between modern humans and archaics, as described below.

We then inspected the 28,491 random SNVs to see whether the derived alleles are shared with any of the four high coverage archaic genomes (Altai Neanderthal, Vindija Neanderthal, Chagyrskaya Neanderthal, and the Denisovan). We found that 4,616 SNVs (16.2%) had their derived allele shared (either homo-or heterozygously) with at least one of the archaics. Since we are only interested in polymorphisms older than 700,000 years, we want to focus only on SNVs where the derived allele is shared with archaics by *common descent*. We thus want to exclude the SNVs where the derived allele emerged independently (recurrence) in humans and archaics. For each of the 4,616 SNVs with shared derived alleles in archaics, we tested whether any of the linked derived alleles is also present in any of the same archaics which carry the derived allele for the random SNV itself. If any of the archaics contain both the derived allele of the random SNV along with at least one linked derived allele, we classify that random SNV as ‘shared by common descent.’ If any random SNV has a derived allele shared with archaics but not a linked derived allele, we classify the random SNV as ‘recurrent.’ This approach yielded 3,894 SNVs (13.7%) wherein the derived allele is shared with at least one of the archaics by common descent.

In order to investigate whether this percentage (13.7%) of shared derived alleles (700,000 years old polymorphisms) is significantly higher than the neutral expectation, we wanted to calculate the same percentage for a set of neutrally simulated SNVs. We used the program *ms* (Hudson, 2002) to neutrally simulate a set of 1000 (independent, and therefore freely recombining) variants (this set of variants was simulated 10,000 times), using the parameters outlined in (Bergström et al., 2021; Gravel et al., 2011; Mafessoni et al., 2020; Meyer et al., 2012; Prüfer et al., 2014). In particular, we assumed that the modern human lineage diverged from archaics ∼700,000 years ago, Denisovans diverged from Neanderthals ∼400,000 years ago, the Altai Neanderthal lineage separated from the Vindija-Chagyrskaya lineage ∼130,000 years ago, and the Vindija lineage separated from the Chagyrskaya lineage ∼90,000 years ago. Additionally, we assumed that the effective size of the population ancestral to archaics and humans, as well as the effective population size of the lineage leading to the modern Yoruba was 14,474. The effective population size for all branches of the archaic lineage was 1,000. These parameters are summarized in **figure S4**. Moreover, the generation time was assumed to be 29 years (Fenner, 2005; Langergraber et al., 2012; Li & Durbin, 2011). We calculated the distribution, based on 10000 runs of simulation, of the percentage of simulated SNVs in Yoruba wherein the derived alleles are shared (by common descent) with archaics. We found that the entire distribution of sharing in our simulated dataset (median=7.8%) lies to the left of the observed sharing of 13.7%. This suggests humans carry significantly more polymorphisms that are older than 700,000 years than expected under neutrality.

Next, we performed another set of neutral simulations, this time with structure introduced in the population ancestral to the archaic hominins and AMH. This too was done using Hudson’s *ms*. The same parameters as the initial neutral simulation (described above, **figure S4**) were used with the following modifications. Going pastward from 700,000 years ago, we divided the ancestral population into three latent subgroups. The effective population size for each of these subgroups was set to 10,000. We defined *m*_*i,j*_ as the fraction of subgroup *i* that is formed by the migrants of subgroup *j* in each generation, where *i ≠ j* and *i,j* ∈ *{1,2,3}*. For all *i and j*, where *i ≠ j*, we set *m*_*i,j*_ = *m*. The program *ms* takes this parameter in the form of *M* = 4N*m* (where N = 10,000 is the effective population size of each subgroup). We performed simulations for 10,000 different values of *M* chosen uniformly on the log scale from the range *(0.01,100)*. This is akin to running simulations using 10,000 different values of *m* in the range *(0.25 × 10*^*-7*^, *0.25 × 10*^*-2*^). For each *m*, 1000 variants were simulated. Thus, for each *m*, we calculated the percentage of variants in Yoruba (with allele-count >1) wherein the derived allele was shared with the archaic hominins. The proportion of allele-sharing in simulations equaled or exceeded the proportion (13.7%) observed in real-life at approximately *m* ≤ 0.0075%.

### Age of deletions and allele frequency trajectories

We estimate the ages of the deletions in the *deletion dataset* using two methods: 1) Human Genome Dating database (https://human.genome.dating/download/index); and 2) RELATE (Speidel et al., 2019).

The Human Genome Dating database (https://human.genome.dating/download/index) hosts age estimates for over 45 million single nucleotide variants (SNVs) (Albers & McVean, 2020). This database reports multiple age estimates for each SNV. We used the median age estimate calculated using the joint clock. Since this database only includes age estimates for SNVs (and not for deletions), we could only date a deletion if the dating database contained the age estimate for at least one of the variants linked (r^2 > 0.9) to the deletion. If age estimates were available for only one linked variant, the same age estimate was assigned to the deletion. If age estimates were available for more than one variant linked to the deletion, we used the highest age estimate, which may be inaccurate in certain cases.

Relate is a method that estimates genome-wide genealogies and can be used to infer the age of a variant (Speidel et al., 2019). We used Relate to infer the ages of the deletions in the *deletion dataset*. To this end, we used previously inferred genome-wide genealogies for samples of the SGDP dataset (Mallick et al., 2016; Speidel et al., 2021), available from https://www.dropbox.com/sh/2gjyxe3kqzh932o/AAAQcipCHnySgEB873t9EQjNa?dl=0. For each deletion, we restricted to tagging SNVs where the derived allele was tagging the deletion at an r^2 exceeding 0.9 and calculated the mean age of such SNVs to date each deletion.

To quantify allele frequency changes, we computed the ratio of lineages carrying the derived allele by the total number of lineages remaining at 5,000 years and 50,000 years before present, but only if the number of lineages remaining at 50,000 years exceeded 10% of the present-day sample size. We then standardized the allele frequency change stratified by present-day allele frequency, by calculating the mean and standard deviation given present-day frequency. Finally, we squared this standardized allele frequency change to obtain our statistic χ^2^, which is expected to have a Chi-squared distribution with one degree of freedom under neutrality, and smaller values for more stable trajectories. This approach was inspired by (Edge & Coop, 2019), who used a similar approach to quantify polygenic positive selection using genealogies.

### Beta measure

We used a recent and robust measure of balancing selection (Siewert & Voight, 2017), ***β***2, to investigate whether ancient deletion are enriched for targets of balancing selection. For a given core allele, ***β***2 measures the weighted average of the number of flanking derived variants, where weights are the similarity in frequency between the core allele and the flanking variants. Suppose that a deletion polymorphism is subject to balancing selection. This deletion will sweep up in frequency until it reaches an equilibrium frequency. Let us refer to the set of all haplotypes (or chromosomes, if recombination is ignored) carrying this deletion as the deleted-class. Single nucleotide variants will have emerged on various haplotypes within the deleted-class since the emergence of the deletion. Some of these single nucleotide variants will drift up in frequency. However, since these variants are linked to the balanced deletion, they will not drift up to fixation since that will require the balanced deletion to reach fixation too. Therefore, the maximum frequency to which they can drift up is the equilibrium frequency of the balanced deletion. We call this phenomenon “Goldilocks drift” since variants are drifting up to the frequency that is “just right” (equilibrium frequency of the balanced deletion). Hence, if a deletion has evolved under balancing selection, the SNVs in its vicinity will have evolved under this Goldilocks drift. Therefore, we should expect an excess of SNVs around the balanced deletion at a similar frequency as that of the balanced deletion itself. A high ***β***2 for a variant is indicative of balancing selection.

In our study, we calculated a variant of the ***β***2 statistic called std***β***2 for the deletions in the deletion dataset. We did this for the CEU, CHB, and YRI population separately. The std***β***2 scores for SNVs are publicly available (https://github.com/ksiewert/BetaScan) for the CEU, CHB, and YRI populations. For each deletion in each of these populations, we obtained the std***β***2 values for the linked variants whenever they were available. When the std***β***2 values were available for more than one linked-variant, we used two approaches to estimate the std***β***2 for the deletions. In the first approach, we used the highest std***β***2 among the linked variants as the estimate for the std***β***2 for the deletion. We call this BETAMAX. In the second approach, we focused on the std***β***2 values for the SNVs that were linked to the deletion with the highest r^2 value. If multiple SNVs were linked to the deletion with the highest r2 value, we used the std***β***2 value of the SNV that was closest to the deletion among these SNVs as the estimate for std***β***2 for the deletion. We call this BETAPRIME. We performed this process for YRI, CEU, and CHB populations separately to arrive at std***β***2 estimates for deletions in our deletion dataset in each of these three populations. Using both BETAMAX and BETAPRIME gave us similar trends across populations.

### Simulations to identify signatures of overdominance

We set out to identify signatures associated with a locus that has evolved under overdominance. Methodologically, we approached the problem of separating overdominance from neutrality on two fronts: (i) The trajectory of the allele-frequency of the mutation conditioning on the age of the mutation, and (ii) the patterns of neutral polymorphisms around the so-called focal mutation which has evolved either under balancing selection or neutrally (with the same age of the mutation). Given that a mutation under overdominance (heterozygote advantage) may be at intermediate frequency, we also (iii) studied simple coalescent simulations conditioning on the existence of at least one SNP within a certain frequency range (e.g. 50%-60%). The age of a mutation is crucial for the study of balancing selection. We considered two values for the age of a mutation: (i) 10,000 generation and (ii) 40,000 generations old. In the first case, the age of the mutation corresponds to 10,000 × 29 years = 290,000 years old mutation. In the second case, the mutation is 1,160,000 years old. Conditioning on the age of the mutation, we generated trajectories of the mutation, i.e., the frequency of the mutation at each time point from its onset until the present-day. For the overdominance scenario, We used the software *trajdemognpops*, implemented using tools from *ms* and *mssel* (kindly provided by R.R. Hudson), to generate trajectories of a mutation under balancing selection. The dominance coefficient is characterized by a large value (here, *h = 10*) in order to assign a benefit for the heterozygote. Thus, the fitness for genotypes at a biallelic locus is by

***Aa***: (1.+*sh*), where *s* is the selection coefficient and *h* the dominance coefficient. Here, *s =* 0.005 and *h* = 10, i.e., the heterozygote Aa has a fitness value 1.05. If we had considered codominance (as opposed to overdominance), we would have set *h* = 0.5.

***AA***: 1 + *s*, thus, the fitness for the AA is 1.005

***aa***: 1.

Given the trajectory of a mutation, a population is split in two kinds of genotypes. The haplotypes carrying the mutation (or the derived allele) and the haplotypes that carry the ancestral allele. Each *neutral* (i.e., passenger mutation) can change ‘population’ (genetic background) by recombination. Therefore, the coalescent in this case is described as a structured coalescent of two populations (a population with the derived allele and another population with the ancestral allele) that communicate between themselves via recombination.

The size of each population is determined at each time point by the trajectory of the derived allele.

In order to understand the effect of the age of the allele (and also to test the mssel code for correctness), we performed coalescent simulations (using Hudson’s ms) conditioning on the presence of at least one SNP at frequency within a given range. Since we are interested in the mutations that are approximately at 50% frequency in the population, we conditioned on the presence of derived mutations in the sample in 22-28 (out of 50) haplotypes. This set of simulations are called *pseCoal*. For each simulated dataset, we calculated the relevant population genetics statistics available through Comus (Papadantonakis et al., 2016) including number of segregating sites, theta, pi, Tajima’s D, ZnS, Fay and Wu’s H, dvk, dvh. Then, we conducted a summary of all these statistics using PCA. The goal is to understand whether the different scenarios can get separated by using polymorphic patterns.

### Conceptual and methodological concerns

#### Human polymorphisms wherein derived allele is shared with archaics are older than 700,000 years

Humans and archaics are estimated to have diverged around ∼700,000 years ago. Therefore, if a human polymorphism has been maintained for more than ∼700,000 years, it was also present in the common ancestral population of human and archaic lineages. It follows that if a polymorphism (the presence of both ancestral and derived alleles in a population) is present in modern humans and archaics, then (barring recurrence and introgression), by parsimony, the polymorphism was also present in their common ancestral population (**Figure 1a**). Thus, a polymorphism that is shared by common descent between modern humans and archaics, has been maintained for over ∼700,000 years. Moreover, because the ancestral allele is fixed in chimpanzees by definition, human polymorphisms wherein the derived allele is fixed in archaics were also present (in the polymorphic state) in the common ancestral population of archaics and humans. In essence, human polymorphisms for which archaics carry the derived allele (fixed or polymorphic) have been maintained for more than 700,000 years.

#### Why use SNVs (as opposed to deletion polymorphisms) for comparison of the real-life proportion of ancient polymorphisms against neutrally simulated SNV

It is worth explaining why we used SNVs, instead of deletions – the class of variants that we are interested in – for comparing observed sharing with archaics against simulated (under neutral conditions) sharing with archaics. Deletions are not suitable for such a comparison because, in general, they are targets of strong negative selection (Conrad et al., 2006; Kondrashov, 2017; Lin et al., 2015; Lin & Gokcumen, 2019). Thus, negative selection would have purged a large proportion of deletions that emerged in the common ancestral population of humans and archaics. It follows that a smaller proportion of human polymorphic deletions than expected under neutral conditions will be shared by common descent with archaics. Even if balancing selection were inflating the proportion of deletions that are shared with archaics, it would not be observable due to the opposite deflationary effect of negative selection. Since negative selection is not as strong a force in the evolution of SNVs as it is for deletions, this problem would not be as pronounced if we used SNVs instead of deletions for testing this premise. Hence, our choice of SNVs for this analysis. Later in the text, we will expand on the idea that deletions are targets of strong negative selection, with implications for large deletions that remain polymorphic in the human lineage.

#### The vast majority of human deletions are derived relative to chimpanzees

In order to identify deletion polymorphisms that have been maintained in the human lineage for over 700,000 years, we focused on deletions that were present (either polymorphically or fixed) in the four high coverage archaic genomes. This technique would work only for deletions that were derived in humans, relative to chimps. However, variants that have been called as deletions (wherein deletion is the alternative allele) in the 1000 Genomes project may, in fact, be human-specific insertions, such that the reference allele (non-deletion) is derived. To investigate how common this situation is among the 6,635 deletions in our dataset, we lifted over the the coordinates of the deletions from hg19 on to the Chimpanzee reference panTro3 using the LiftOver tool in UCSC Genome Browser (Kent et al., 2002). If the liftover for a deletion fails on account of the window being completely or partially deleted in the Chimpanzee reference, it is indicative of the region being a human-specific insertion. The liftover failed for this reason for only 238 (3.6%) of the 6,635 deletions. Therefore, the vast majority of deletions are, in fact, derived relative to chimps.

**Figure S1.**
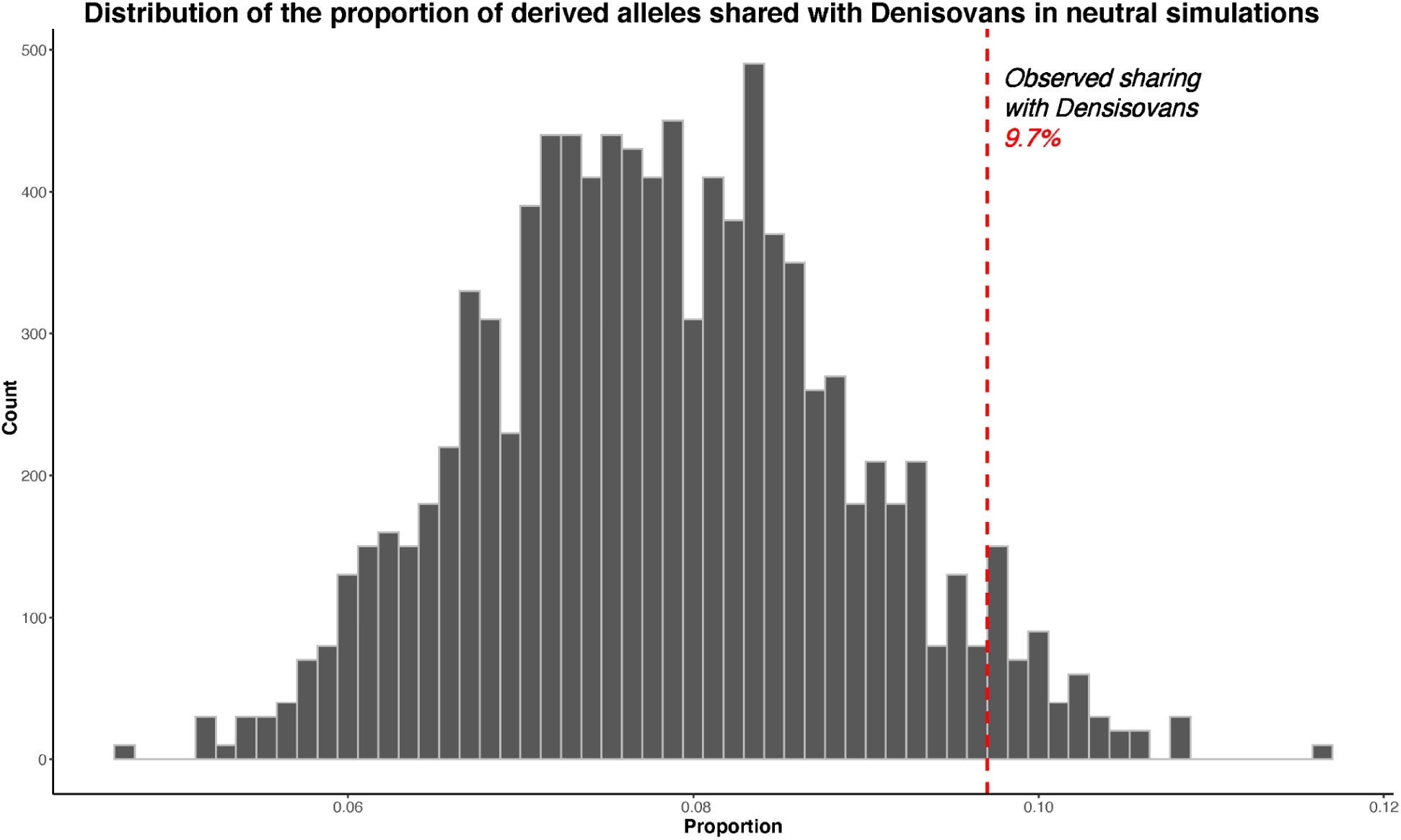
Distribution of the proportion of derived alleles that Yorubans share with the Denisovan genome in 10,000 replicates of neutral simulations. The vertical red line indicates the proportion of derived alleles that Yorubans share with Denisovans in actual data. There is an excess of derived alleles that Yorubans share with Denisovans, relative to neutral expectation.

**Figure S2.**
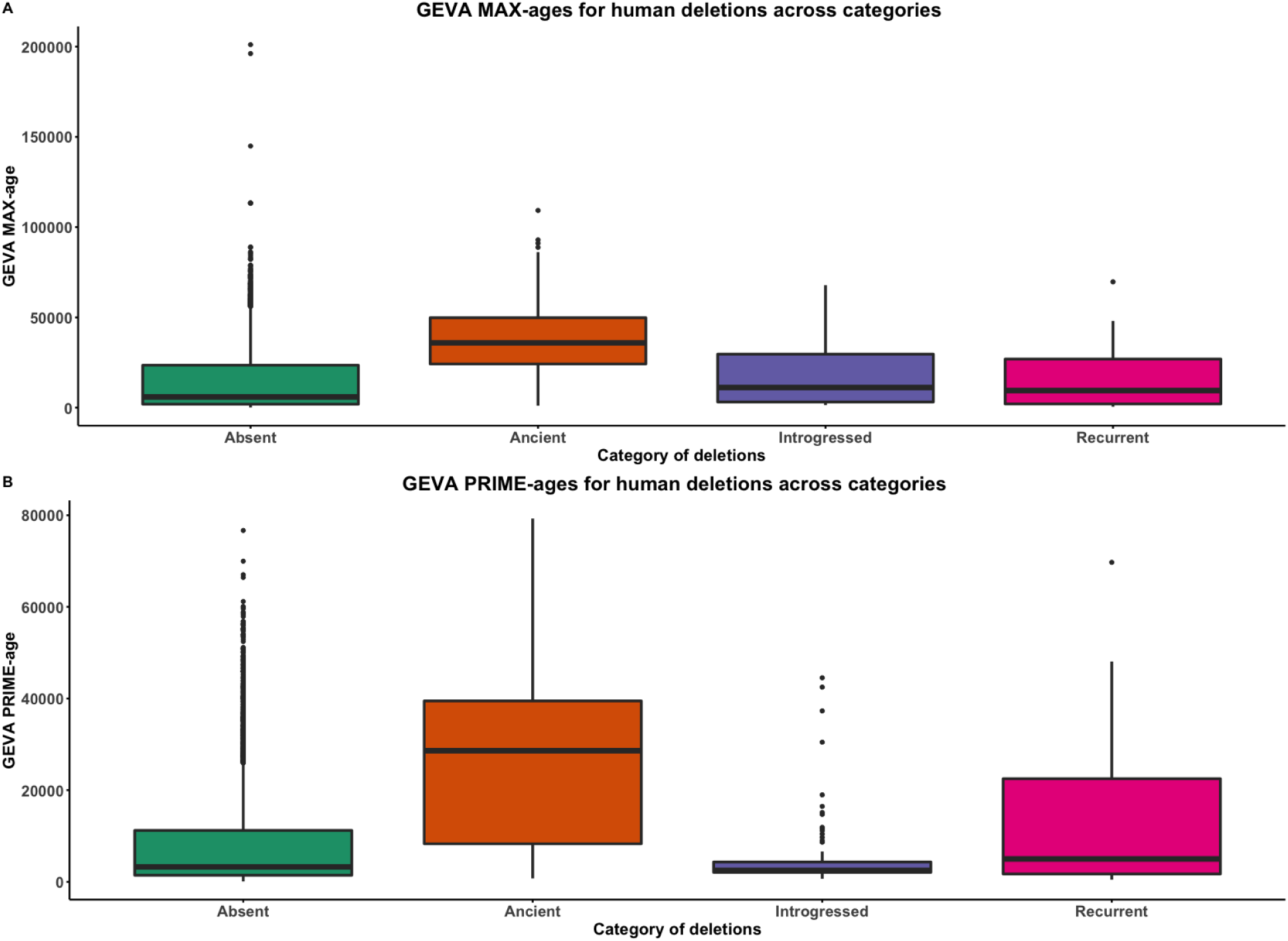
GEVA ages of deletions across categories. Absent denotes human polymorphic deletions that are not present in any of the four high coverage archaic genomes. Introgressed refers to the shared deletions that were introgressed from archaics into modern humans. Recurrent refers to the shared deletions that emerged independently in the modern human and archaic lineages. Ancient refers to the human deletions that are shared with archaics by common descent. **A**. GEVA PRIME-ages. **B**. GEVA MAX-ages. With both GEVA PRIME and GEVA MAX measures, we observe that ancient deletions are significantly older than absent, recurrent, and introgressed deletions. This implies that our pipeline to identify ancient deletions is sound.

**Figure S3.**
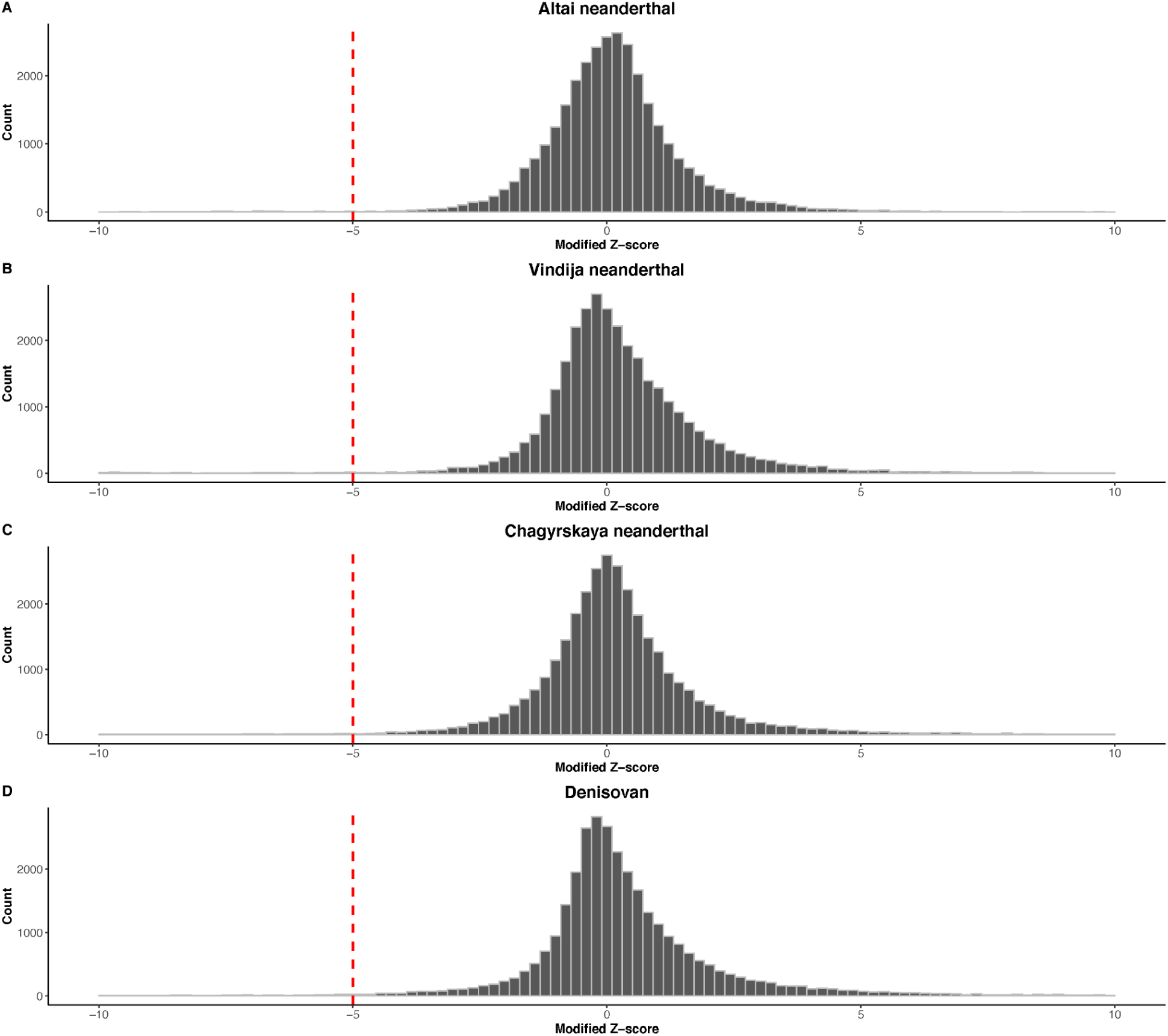
Distribution of the modified Z-score of the read-depth across the 32,154 modern human deletions in the archaic genomes. **A**. Altai neanderthal. **B**. Vindija neanderthal. **C**. Chagyrskaya neanderthal. **D**. Denisovan.

**Figure S4.**
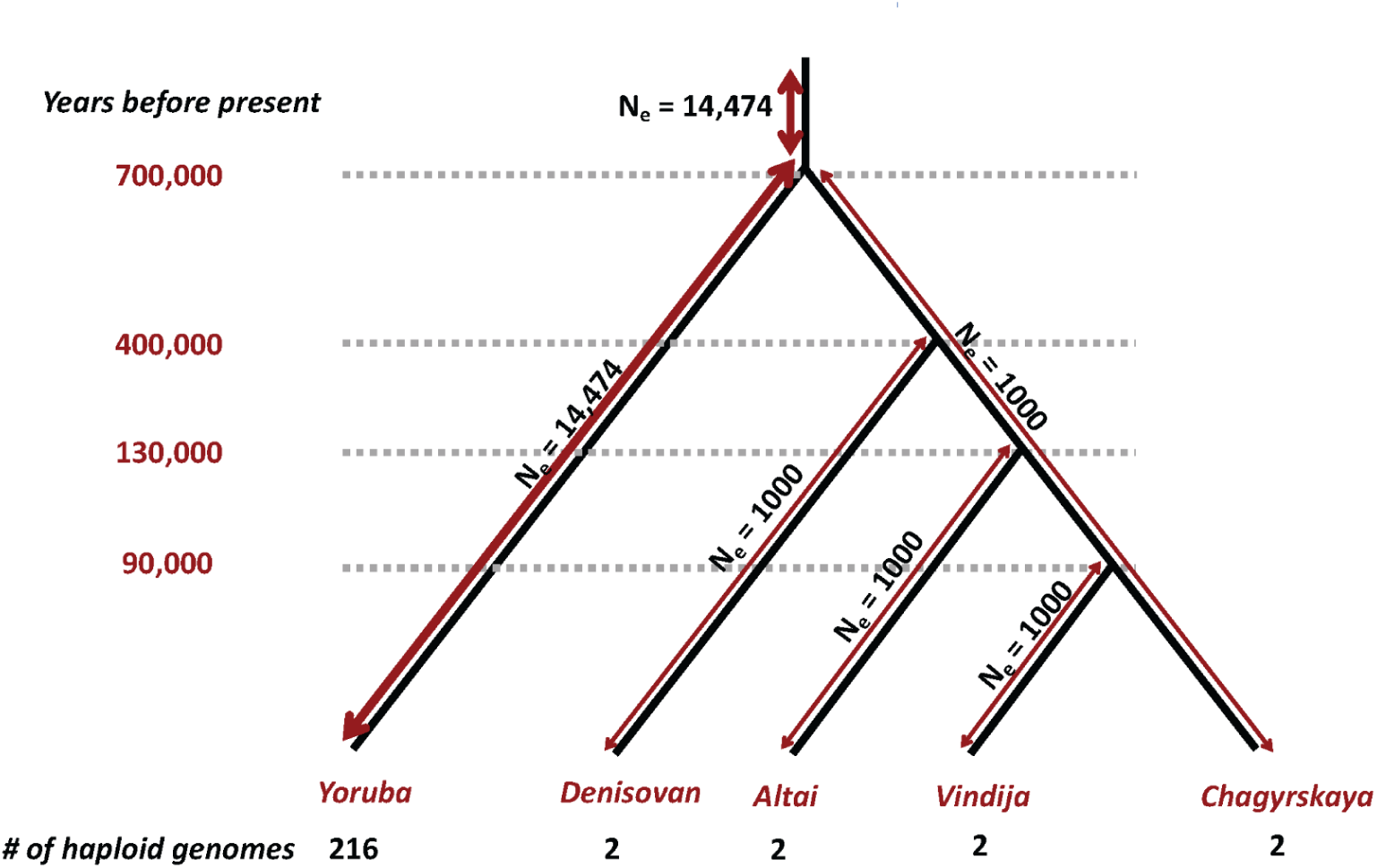
A summary of the parameters used in neutral simulations. The effective population for the YRI lineage all the way up to the common ancestor of archaic hominins and AMHs was assumed to be 14,474. The effective population size of all archaic hominin lineages was assumed to be 1000. The effective population size of the common ancestor of archaic hominins and AMH was assumed to be 14,474. The divergence times of pairs of lineages is mentioned on the left hand side of the horizontal line passing through the point representing the divergence between a pair of lineages.

